# Longitudinal Awake Mouse Brain Imaging Using Functional Ultrasound and Functional Ultrasound Localization Microscopy

**DOI:** 10.64898/2026.05.22.727269

**Authors:** Zhe Huang, Yike Wang, Matthew R. Lowerison, Yue Xu, Bing-Ze Lin, YiRang Shin, Nathiya Vaithiyalingam Chandra Sekaran, Daniel A. Llano, Pengfei Song

## Abstract

Ultrafast ultrasound offers a unique route to cross-scale neurovascular phenotyping by integrating functional ultrasound (fUS), ultrasound localization microscopy (ULM), and functional ULM (fULM). Yet the baseline variability, longitudinal stability, and biological safety of such multimodal imaging in awake animals remain insufficiently defined, limiting its use for detecting subtle disease-associated neurovascular changes. Here, an awake longitudinal fUS-ULM-fULM framework is established and validated in mice over five months. Structural vascularity, microvascular flow velocity, mesoscale hemodynamic responses, and microvascular functional responses are repeatedly quantified in the same animals during monthly imaging sessions. Across all metrics, no significant longitudinal drift is detected (p > 0.60). Structural and flow-derived measures are markedly more reproducible than functional readouts, with within-subject coefficients of variation of 5.1% for mean flow velocity and 7.3% for vascularity, compared with 25.0% for fUS-derived cerebral blood volume responses and 53.2% for fULM-derived microvascular functional responses. Mean flow velocity shows the strongest longitudinal consistency (ICC = 0.70) and the lowest detection threshold. Behavioral testing and GFAP/Iba1 staining further reveal no memory impairment or chronic neuroinflammation. This study defines quantitative baselines, reproducibility limits, and safety evidence for awake cross-scale ultrasound imaging, providing a reference framework for longitudinal neurovascular phenotyping in preclinical disease models.

## 1 Introduction

Many neurological and neurodegenerative disorders, most notably Alzheimer’s Disease and Vascular Cognitive Impairment, are closely associated with abnormalities of the cerebral vasculature, including dysregulated blood flow, vascular remodeling, and impaired neurovascular coupling ^1,2^. Gaining mechanistic insight into these diseases and developing effective therapeutic strategies rely heavily on mouse models, which enable causal and longitudinal investigation of disease processes under well-defined experimental conditions. In this context, advanced neuroimaging is an essential tool for interrogating such abnormalities in the *in vivo* mouse brain. By monitoring the long-term evolution of the cerebral vasculature and characterizing neurovascular homeostasis across physiological and pathological states, such imaging technologies provide a critical foundation for understanding the onset and progression of neurological and neurodegenerative disorders as well as the effectiveness of therapies and interventions.

To meet this requirement, imaging approaches capable of simultaneously capturing large-scale brain territories and fine-scale local regions are essential ^3^. Although existing neuroimaging techniques such as magnetic resonance imaging(MRI) ^4,5^ and optical imaging ^6,7^ have substantially advanced our understanding of the brain, they face inherent limitations in the context of imaging territory and spatial resolution. MRI provides whole-brain coverage but lacks the spatial resolution and specificity to resolve microscopic vasculature and micro-circuitry-level brain activity ^4,5^. In contrast, optical imaging provides exquisite microscopic resolution but suffers from limited imaging depth ^6,7^, which confines its utility to superficial brain regions. In recent years, ultrasound has rapidly emerged as a promising brain imaging modality ^8^ that bridges the imaging resolution and coverage gap between MRI and optics by using high frequency ultrasound waves that provide both robust imaging depth of penetration and spatial resolution. At present, three major ultrasound brain imaging techniques are used for studying the rodent brain: Functional ultrasound (fUS) is a contrast-free brain blood flow imaging technique that is sensitive to functional hyperemia and provides maps of cerebral blood volume changes (ΔCBV) across brain-wide territories at a mesoscopic spatial resolution ^9^. Ultrasound localization microscopy (ULM) is a contrast-enhanced super-resolution imaging approach that localizes and tracks circulating microbubbles (MBs) in the blood stream to visualize the microvascular architecture and measure their flow ^10,11^. ULM also allows detailed assessments of local and global vessel morphology (e.g., density, tortuosity, intervessel distance) and flow distributions ^12–14^; functional ULM (fULM) is also a contrast-enhanced imaging technique that essentially combines ULM and fUS, which enables microvessel-scale mapping of cerebral blood flow (CBF) dynamics to detect stimulus-locked hemodynamic responses manifested as MB count fluctuations modulated by the underlying neural activity ^15^. Although all three ultrasound imaging modalities rely on blood flow measurements, they differ in functionality and provide complementary information about the brain. For example, both fUS and fULM provide functional mapping of the brain but they differ greatly in spatial (hundreds of microns for fUS, several microns for fULM) and temporal resolution (<1s for fUS and sub-minute timescales for fULM). ULM complements fUS and fULM by serving as a primary structural imaging modality that reveals vascular anatomy and flow distributions. Therefore, integrating fUS, ULM, and fULM into a unified imaging platform will enable comprehensive structural and functional evaluations of the brain across different scales from the whole brain to individual microvessels, providing a powerful tool for studying neurological diseases in rodent models.

Another major advantage of the three ultrasound brain imaging modalities is their suitability for imaging awake or freely moving animals, which is essential for studying brain function under naturalistic conditions without the confounding effects of anesthesia (e.g., isoflurane, which markedly alters CBF ^16^). In addition, ultrasound is inherently well-suited for longitudinal and serial imaging studies because of the lack of ionizing radiation and low cost ^17^. Despite these advantages, however, fUS, ULM, and fULM have primarily been implemented independently in awake or longitudinal imaging paradigms ^14^ and they have not been combined within the same animals in a longitudinal setting to provide an integrated assessment of brain structure and function. Achieving such integration is technically nontrivial as it requires carefully coordinated acquisition protocols, imaging sequences, as well as stable and consistent microbubble administration in awake animals.

To this end, in this study we directly address these challenges by establishing an integrated fUS-ULM-fULM imaging platform for awake mouse brain imaging over time. We conducted a 5-month longitudinal brain imaging study in awake mice, incorporating whisker stimulation during fUS and fULM to probe evoked functional vascular responses, while ULM was used to quantify microvascular structure and flow-related metrics. We also evaluated chronic-window safety and, to confirm that memory function remained intact after repeated imaging, assessed behavior using a passive avoidance task. Our goal is to evaluate the feasibility of combining all three imaging approaches in an awake and longitudinal imaging setting while characterizing the baseline variations (across several months) in the quantitative structural and functional vascular measurements from fUS, ULM, and fULM in a cohort of animals. By quantifying the inherent measurement and physiological variations of each metric, we defined the necessary sensitivity thresholds required to distinguish subtle, early-stage pathological alterations from baseline fluctuations observed in wild-type animals.

## 2 Results

Seven male mice were included in this study. Only male mice were used to reduce potential biological variability associated with sex-dependent vascular and physiological fluctuations, including those related to the estrus cycle ^18^. Each mouse was implanted with a transparent polymethylpentene (PMP) cranial window and allowed to recover for 14 days before imaging. After recovery, the animal was placed in a custom-designed body tube that was securely fixed to the recording table 19(Fig. 1a, 1b). The ultrasound probe was positioned above the brain and coupled with ultrasound gel to ensure optimal acoustic transmission (Fig. 1c). Each mouse underwent longitudinal imaging(Fig. 1d) that included (in the following order) whisker-stimulation-based fUS (Fig. 1e, top row), ULM (Fig. 1e, 2^nd^ and 3^rd^ rows), and whisker-stimulation-based fULM (Fig. 1e, bottom row). Each animal was imaged over a five-month period, with one imaging session conducted per month. Representative ultrasound images from one animal are presented in Fig. 1e. After completion of the longitudinal imaging, the mice underwent behavioral testing and histological staining.

**Fig 1.**
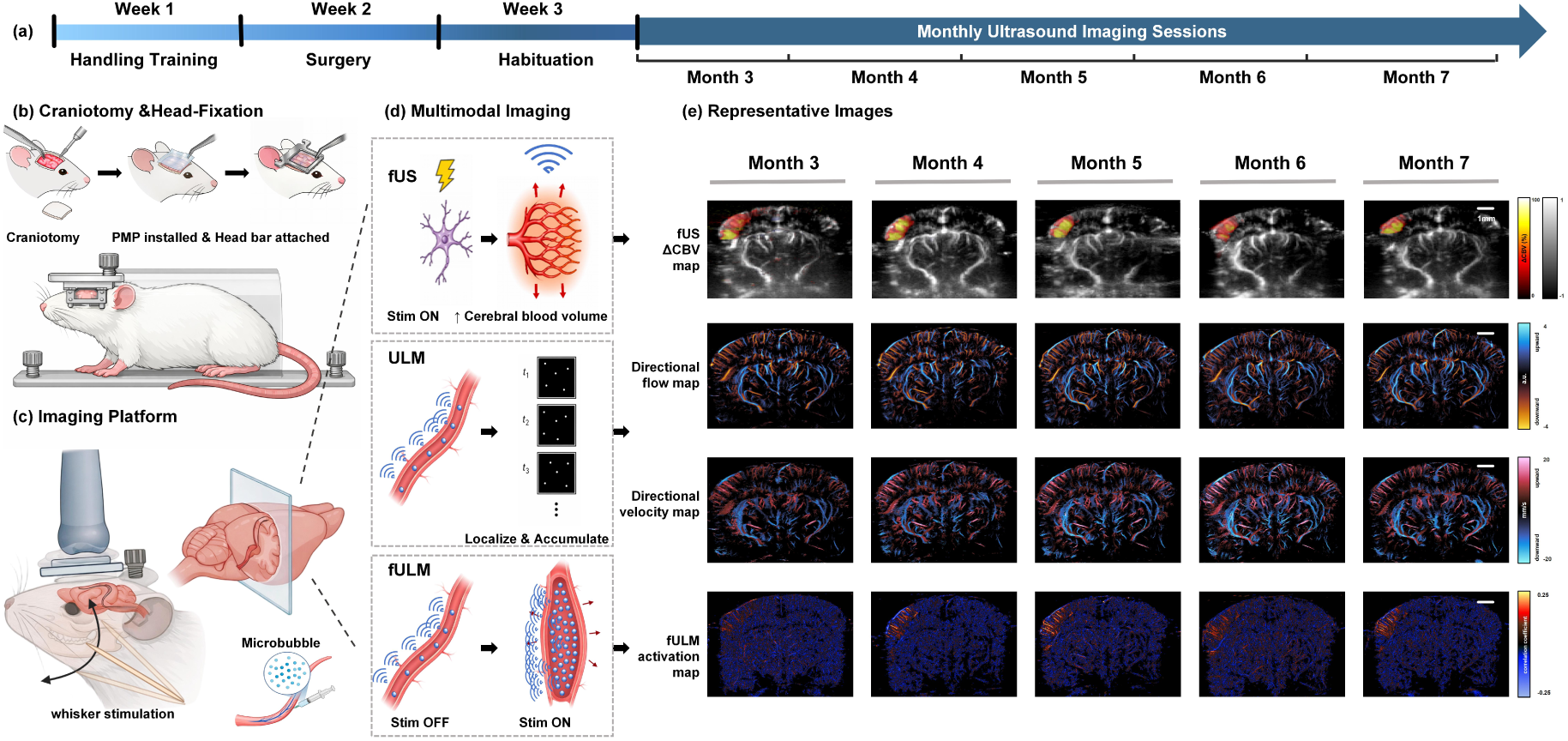
Schematic overview of the longitudinal awake ultrasound imaging workflow. (a) Experimental timeline showing handling training in Week 1, cranial window/head-bar surgery in Week 2, habituation in Week 3, and monthly ultrasound imaging from Month 3 to Month 7. (b–d) Schematic illustrations of the craniotomy and head-fixation preparation, awake imaging platform with whisker stimulation and microbubble injection, and the multimodal imaging strategy integrating fUS, ULM, and fULM. (e) Representative longitudinal images acquired from the same awake mice across five monthly sessions. Whisker-evoked fUS ΔCBV maps show mesoscale hemodynamic responses in the S1 barrel cortex; ULM directional flow and directional velocity maps reveal microvascular architecture and flow dynamics; and fULM activation maps capture stimulus-evoked microvascular functional responses. Created in BioRender. Lab, S. (2026) https://BioRender.com/w1cs5bc

**Fig 2.**
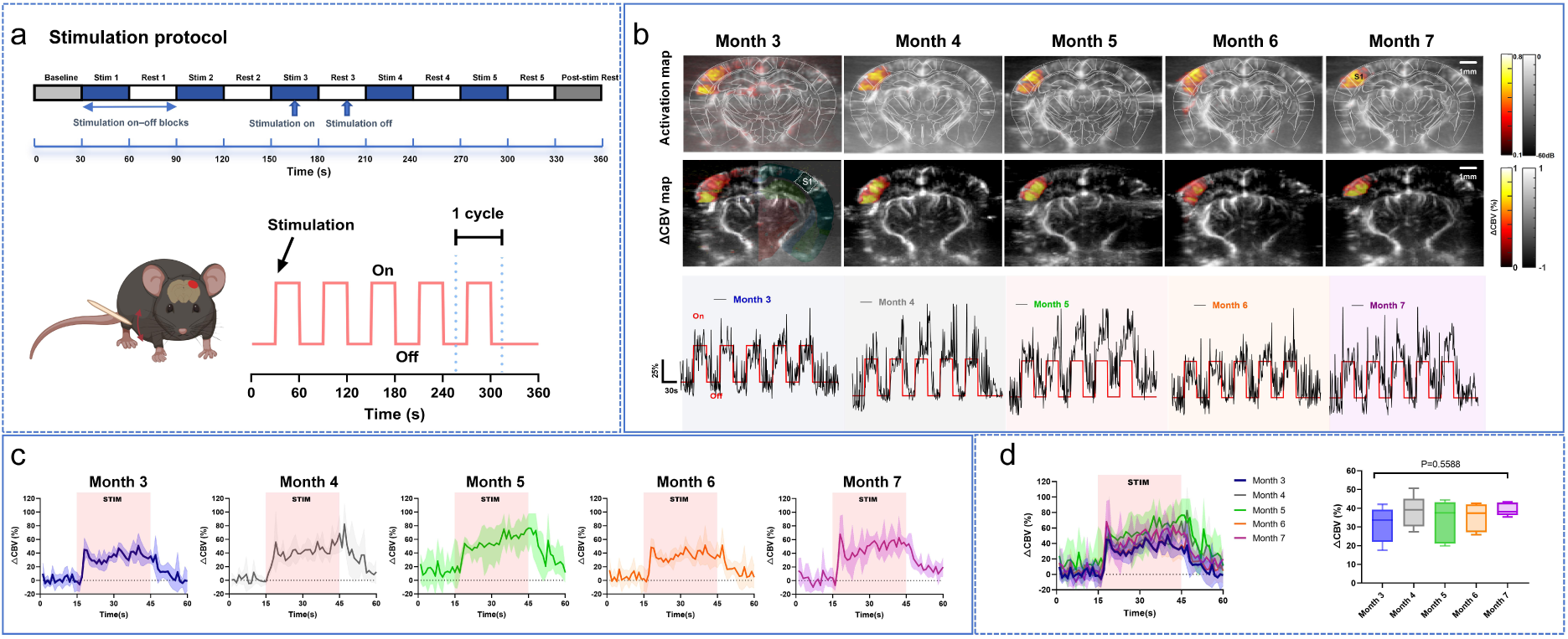
Longitudinal whisker-stimulation-based fUS imaging in awake mice. (a) The fUS imaging session lasted a total of 360 seconds, consisting of 30 seconds of baseline recording, five stimulation on-off blocks (30 seconds of stimulation on followed by 30 seconds of stimulation off per block), and 30 seconds of post-stimulation recovery in the end. (b) Representative fUS activation maps (top row, obtained from correlation between power Doppler signal and stimulus pattern), ΔCBV maps (middle row, obtained from calculating power Doppler signal difference between stimulation on and off), and ΔCBV time series (bottom row) from the same animal imaged longitudinally from 3 to 7 months of age. (c) Averaged ΔCBV time courses within the S1 region, each curve representing one monthly imaging session. For each month, the ΔCBV response was averaged across five stimulation cycles (one cycle defined as the stimulation on/off block shown in panel (b), lower-left). Data are presented as mean (solid lines) ± SEM (shaded area). For this representative mouse, the mean ΔCBV across the five sessions was 42.28%, with a within-mouse CV of 20.5%. (d) Group-level comparison of mean ΔCBV across five monthly sessions(n=7); one-way ANOVA, p=0.2167. Data are shown as box-and-whisker plots (median and IQR). fUS, functional ultrasound; ΔCBV, change in cerebral blood volume; SEM, standard error of the mean; IQR, interquartile range.

**Fig 3.**
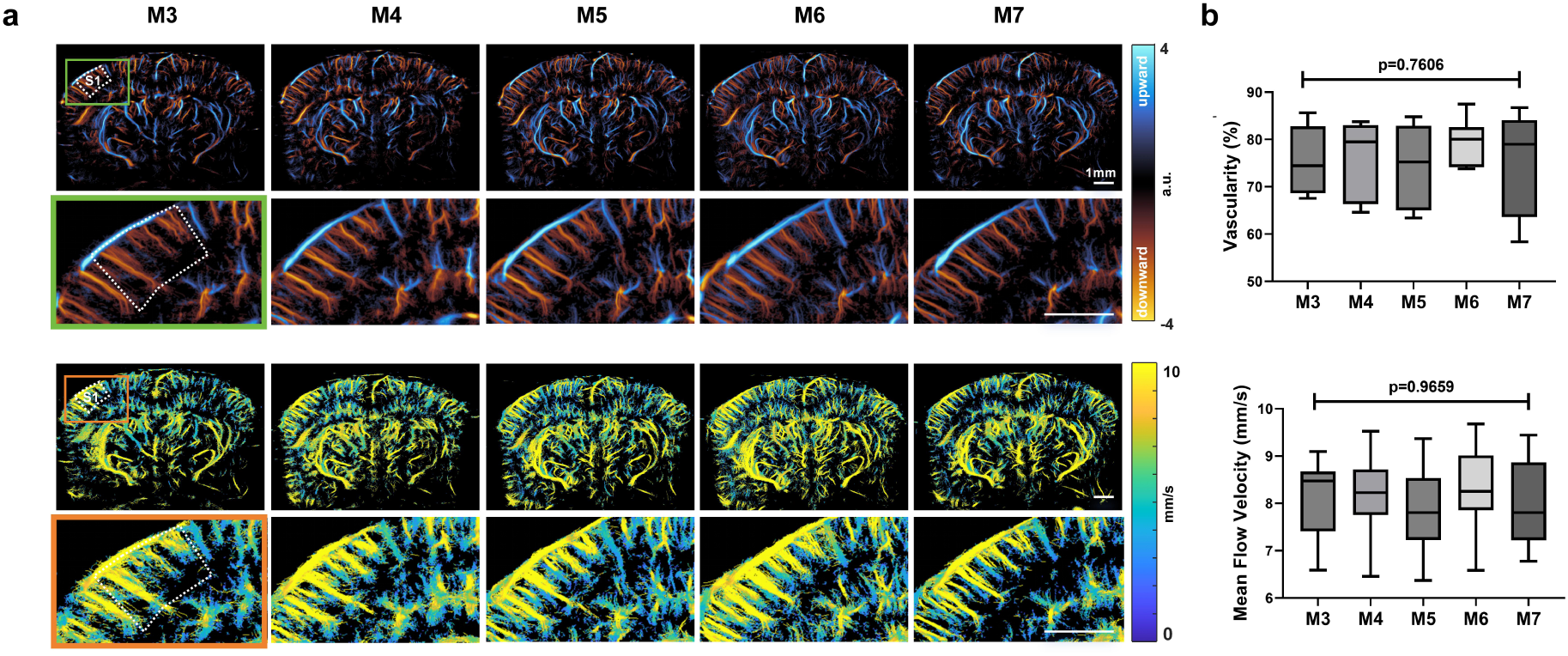
Longitudinal ULM imaging reveals stable microvascular structure and flow velocity. (a) Directional flow maps specifically within the S1 region illustrating arterial (red, downward flow) and venous (blue, upward flow) components from 3 to 7 months. Flow velocity maps acquired from 3 to 7 months. (b) Group-level analysis of vascularity and flow velocity within the S1 region (n = 7). Data are presented as box-and-whisker plots showing the median and IQR. ULM, ultrasound localization microscopy; S1, primary somatosensory cortex; IQR, interquartile range.

### fUS Consistently Measures Mesoscale Brain Functional Responses to Whisker Stimulation

We characterized whisker stimulation-evoked functional hyperemia in awake mice from 3 to 7 months of age using fUS. Across all longitudinal imaging sessions, whisker stimulation consistently elicited robust hemodynamic activation in the contralateral primary somatosensory cortex (S1), with significant ΔCBV response consistently localized to the barrel field (Fig. 2a,b,c). No stimulus-locked ΔCBV changes were observed in ipsilateral S1 or non-sensory cortical regions, which confirms the spatial specificity of the response. Using mouse brain atlas ^20^, stimulus-evoked functional activation remained localized to a similar cortical region over the five-month imaging period. To quantify within-mouse longitudinal spatial reproducibility, we used month 3 as the reference session and calculated the Jaccard index between the binarized activation map at month 3 and those at months 4, 5, 6, and 7. Across the seven mice, the mean Jaccard indices were 0.78 ± 0.11, 0.82 ± 0.09, 0.85 ± 0.13, and 0.81 ± 0.15, respectively, supporting stable spatial overlap of the evoked response across sessions. Quantitative analysis of ΔCBV time courses shows that the magnitude of the whisker stimulation responses exhibited no significant month-to-month variations (one-way ANOVA, p = 0.2167; Fig. 2d), with response amplitudes maintained within a range of 29-39%, with an overall coefficient of variation (CV) of 24.5% across all animals and sessions. fUS ΔCBV had a within-mouse CV of 25.0%, an intraclass correlation coefficient (ICC) of 0.48, a repeatability coefficient (RC) of 24.8%, and no significant longitudinal trend (linear mixed-effects models, trend p = 0.688; Table 1), indicating moderate session-to-session variability without evidence of systematic drift over time. Paired TOST analysis supported equivalence between Month 3 and Month 7, with a 90% confidence interval of 0.004% to 16.28% fully contained within the predefined equivalence margin of ±24.8%. Together, these findings indicate that fUS can repeatedly detect whisker stimulus-evoked functional hyperemia in awake mice over multiple months, with reproducible spatial localization across sessions and no evidence of systematic drift in the mean response over time.

**Table 1.**
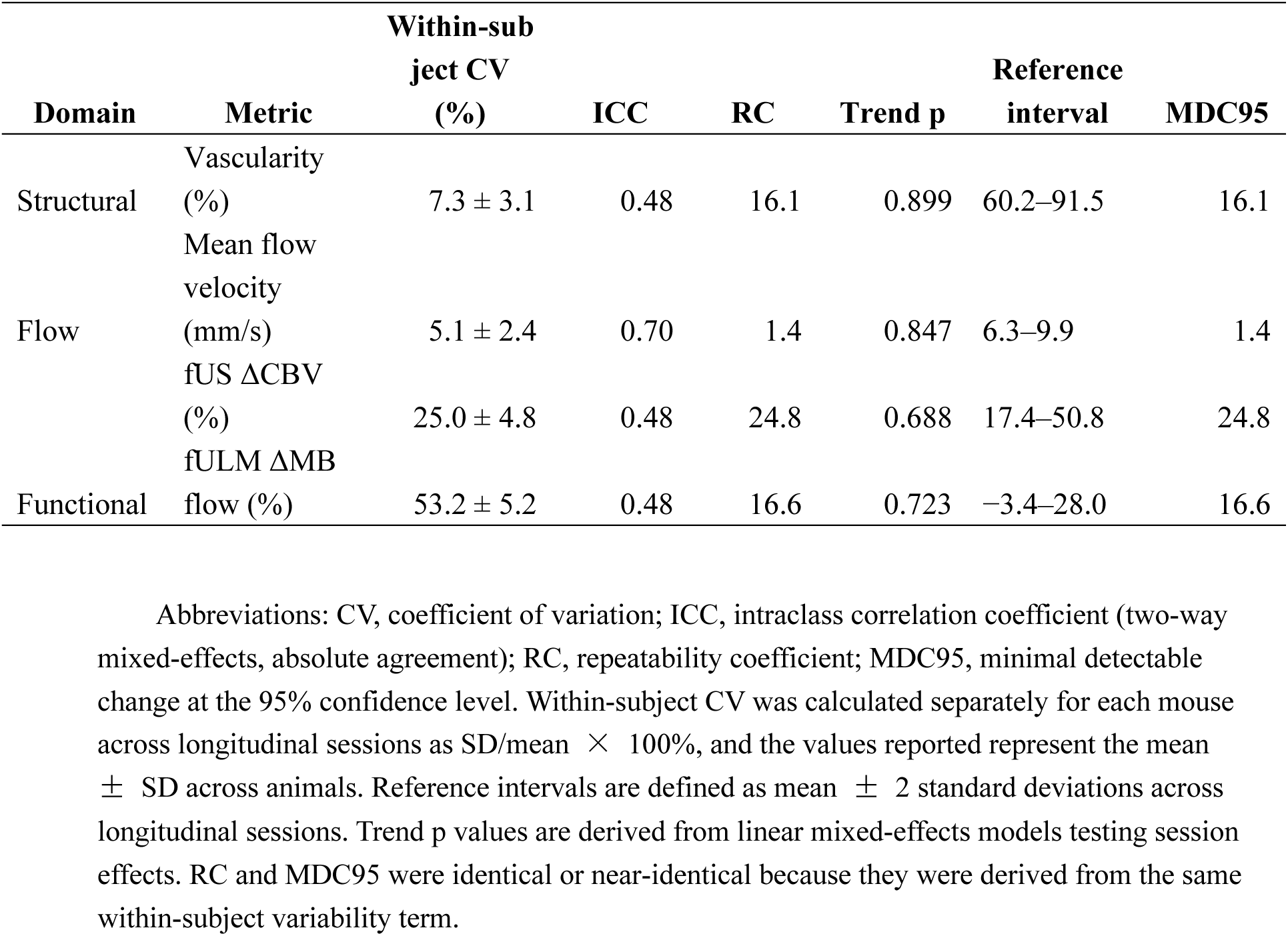
Longitudinal variability, reproducibility, and detection thresholds of multimodal ultrasound metrics. Abbreviations: CV, coefficient of variation; ICC, intraclass correlation coefficient (two-way mixed-effects, absolute agreement); RC, repeatability coefficient; MDC95, minimal detectable change at the 95% confidence level. Within-subject CV was calculated separately for each mouse across longitudinal sessions as SD/mean × 100%, and the values reported represent the mean ± SD across animals. Reference intervals are defined as mean ± 2 standard deviations across longitudinal sessions. Trend p values are derived from linear mixed-effects models testing session effects. RC and MDC95 were identical or near-identical because they were derived from the same within-subject variability term.

### ULM Indicates Stable Microvascular Architecture and Flow Velocity

Figure 3 presents the ULM imaging results across five imaging sessions throughout 5 months. Within the S1 cortex (white dotted ROI in Fig. 3a), ULM-derived vascularity remained stable across all time points, with no significant month-to-month differences observed at the group level (one-way ANOVA, p = 0.7607; Fig. 3b). Vascularity values remained within a narrow range (74.0%-79.0%) across sessions, with an overall CV of 10.0%, indicating stable microvascular structural organization over time. Mean cerebral microvascular flow velocity within the S1 cortex also remained stable over time (one-way ANOVA, p = 0.9659; Fig. 3b). The velocity fluctuated within a narrow range (7.9-8.2 mm/s; overall CV = 11.1%). Consistent with these group-level observations, longitudinal modeling further supported the stability of both ULM-derived metrics. Linear mixed-effects analysis showed no significant longitudinal trend for either mean flow velocity or vascularity (trend p = 0.847 and 0.899, respectively), together with relatively low within-mouse variability (5.1 ± 2.4% and 7.3 ± 3.1%) and ICC values of 0.70 and 0.48 (Table 1). Paired TOST analysis further supported equivalence between Month 3 and Month 7 for both metrics, with 90% confidence intervals fully contained within the predefined MDC95-based margins (mean flow velocity: −0.849 to 0.694 mm/s within ±1.4 mm/s; vascularity: −8.53% to 4.59% within ± 16.1%). Across ULM-derived measures, variance decomposition indicated that within-subject variability accounted for a minority of total variance, with inter-animal differences predominating. Some residual variation may reflect vessel blurring caused by residual tissue motion during awake imaging.

### fULM Indicates Stable Microvessel-level Hemodynamic Responses to Whisker Stimulation

Figure 4 shows the representative activation maps generated by fULM which reveal spatially localized microvascular responses. Within the S1 region, fULM-derived ΔMB flow (%) time courses exhibited synchronized blood flow increases within the stimulation window (15-45 s; Fig. 4a,b,c,e). Because fULM measures blood flow fluctuations based on microbubble counts, which is different from fUS, the ΔMB flow (%) measurements range from 9-17% with an overall CV of 63.9% across all animals and sessions, indicating greater variability than fUS. fULM analysis across all five monthly sessions (3-7 months) demonstrated no significant longitudinal changes in mean ΔMB flow (%) responses (one-way ANOVA, p = 0.6080; Fig. 4e). Similarly, the activation measurements derived from correlation between microbubble fluctuation and the stimulus pattern remained stable across longitudinal sessions (one-way ANOVA, p = 0.9747; Fig. 4f). fULM-derived ΔMB flow (%) also showed no significant longitudinal trend by linear mixed-effects analysis (trend p = 0.723), but exhibited the greatest within-mouse variability among all readouts (53.2 ± 5.2%), with an ICC of 0.48 (Table 1). Paired TOST analysis nevertheless supported equivalence between Month 3 and Month 7, with a 90% confidence interval of − 8.12% to 4.20% fully contained within the predefined equivalence margin of ±16.6%. Together, these findings indicate that fULM can be reproducibly and longitudinally implemented in an awake imaging setup.

**Fig 4.**
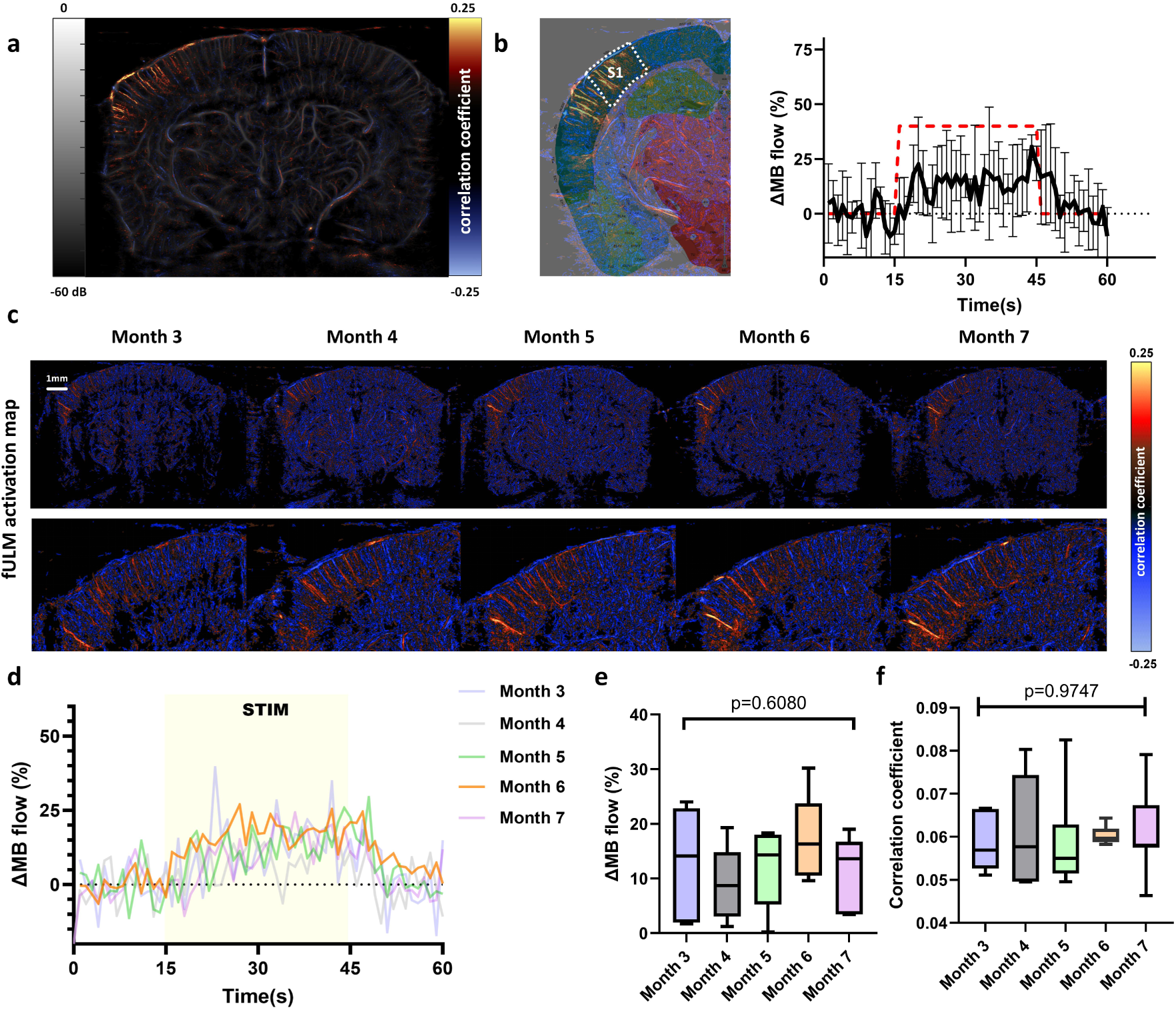
Longitudinal fULM imaging and microvessel-level stimulus-evoked hemodynamic responses. Representative fULM activation maps during right-whisker stimulation, showing localized responses in the contralateral S1 cortex. (b) Averaged ΔMB flow (%) time course within the S1 region from the same animal as in (a), derived from five stimulation cycles (red dashed lines indicate the 15-45 s stimulation period; data are mean ± SEM). (c) Longitudinal fULM activation maps from the same mouse illustrating consistent microvessel-level activation across months. (d) Averaged ΔMB flow (%) time course within the S1 region derived from five stimulation cycles of 5 months. (e, f) Group-level analyses (n = 7) showing stable mean ΔMB flow (%) and flow-stimulus correlation coefficients across five months. Data are presented as box-and-whisker plots showing the median and IQR. fUS, functional ultrasound; fULM, functional ultrasound localization microscopy; ΔCBV, change in cerebral blood volume.

### Longitudinal Baseline Variability, Reproducibility, and Detection Thresholds

Table 1 summarizes the longitudinal baseline variability and reproducibility statistics of multimodal ultrasound measurements acquired over the five monthly awake imaging sessions. No metric showed significant longitudinal trends, with all *p* values above 0.60, which indicates the absence of systematic time-dependent differences (Table 1). Month-to-month variability was lower for structural and flow metrics than for functional responses, with within-subject CVs of 5.1% for mean flow velocity, 7.3% for vascularity, compared with 25.0% for fUS-ΔCBV and 53.2% for fULM-ΔMB flow (%) (Table 1). Mean flow velocity showed the strongest between-animal consistency relative to within-subject variation (ICC = 0.70) (Table 1).

Table 1 also summarizes the detection thresholds for each quantitative metric, providing practical reference values for future studies designed to detect biological differences using the fUS, ULM, or fULM. Mean flow velocity showed the lowest detection threshold, requiring a change of 1.4 mm/s to exceed baseline variability. Vascularity required a change of 16.1% to exceed baseline variability. Functional readouts required changes of 24.8% for fUS-ΔCBV and 16.6% for fULM-ΔMB flow (%) for detectable differences. Together, these results indicate that mean flow velocity was the most consistent longitudinal readout, whereas functional responses were more variable and therefore required larger effect sizes for reliable detection.

### Chronic PMP Cranial Window Maintains Long-Term Cortical Biocompatibility

To investigate whether the long-term cranial window implantation induced chronic inflammation to the brain, we performed immunofluorescence staining for GFAP (astrocytes) and Iba1 (microglia) after the five-month imaging protocol (Supplementary Fig 1a). Quantitative analysis demonstrated that GFAP and Iba1 expression levels beneath the PMP window were comparable to those in the non-window-covered cortex, indicating no significant glial activation (Supplementary Fig 1b). Consistent with this observation, we found no morphological features of reactive astrogliosis or microglial aggregation, which are hallmarks of chronic neuroinflammation or foreign body responses. Fluorescence intensity profiles for both markers were consistent across animals, and no focal accumulation of GFAP or Iba1 was detected beneath the window. These results indicate that the surgical implantation of the PMP window, together with repeated head fixation and longitudinal imaging, does not induce sustained cortical inflammation or scarring. To evaluate whether cognitive function was affected after 5 months of repeated imaging, we assessed behavior using a passive avoidance task. Mice exhibited prolonged step-through latencies during the retention test (Supplementary Fig. 1c), consistent with preserved avoidance memory.

## 3 Discussion

In this study, we established an integrated multiscale ultrasound imaging framework combining fUS, ULM, and fULM for awake longitudinal mouse brain imaging. Using a chronically stable PMP cranial window, we demonstrated serial multimodal acquisition in the same animals over five months and quantified baseline variability across mesoscopic hemodynamic and microvessel-scale structural/flow metrics, providing sensitivity benchmarks for distinguishing disease-related deviations from baseline fluctuations.

Using fUS, we detected robust whisker-evoked hemodynamic responses in the contralateral barrel cortex of awake mice. Activation maps were used to extract ΔCBV time courses, which showed an onset within the first few second followed by a slower rise, consistent with the expected hemodynamic response dynamics. Compared with prior reports in anesthetized mice (relative CBV changes of ∼10.5 ± 3.7% at 2 months and ∼8.7 ± 2.7% at 10 months ^21^), responses in awake mice were larger (ΔCBV 29–39% from 3–7 months), which is consistent with the known influence of anesthesia on neurovascular coupling. Our study showed that awake imaging is associated with greater longitudinal variability, and our statistical analysis showed that fUS-ΔCBV changes need to exceed 24.8% to be detectable over baseline variability. These results provide a practical benchmark for interpreting biologically meaningful differences in future awake imaging studies using fUS. We did not observe clear ventral posteromedial nucleus activation under the present stimulation and imaging conditions. Although whisker-evoked VPM responses have been reported in previous ultrasound studies ^22^, their detectability may vary depending on imaging modality, sensitivity, and the experimental configuration, particularly in the awake state. In contrast, cortical activation patterns can appear spatially broader, extending beyond the S1 barrel core into adjacent cortex, consistent with thalamocortical and intracortical propagation as well as vascular integration across shared arteriolar territories ^23,24^.

ULM enables super-resolution ultrasound imaging of the cerebral microvasculature, allowing visualization of microvessel networks beyond the resolution of conventional ultrasound ^25^. In healthy awake mice, ULM-derived metrics (vascularity and mean flow velocity) remained stable over five months, providing a quantitative baseline for longitudinal studies and for detecting deviations induced by aging or disease. To our knowledge, this work represents the first longitudinal demonstration of ULM imaging stability in awake mice. Within the S1 region, ULM flow velocity showed the strongest reproducibility and lowest longitudinal variability (CV = 5.1%), with a minimal detectable change of 1.4 mm/s, while vascularity also exhibited low variability (CV = 7.3%) and required a 16.1% change to exceed baseline variability. These findings support the longitudinal stability of ULM and demonstrate that our awake longitudinal imaging procedure enables reliable repeated structural and flow quantification over months in awake animals.

fULM extends ULM by enabling microvessel-scale mapping of stimulus-evoked flow dynamics ^13,26^. In the present study, fULM resolved localized microvascular responses to whisker stimulation that are not accessible with fUS imaging alone. Although fULM-derived functional metrics showed greater within-subject and session-to-session variations than mesoscopic fUS measures, they remained reproducible at the cohort level, with no evidence of systematic longitudinal drift across repeated sessions. These findings support fULM as a reproducible tool for establishing baseline microvascular functional references and for future longitudinal studies of neurovascular coupling alterations in awake mice.

Our study has several limitations. First, the head holder and residual skull bone introduced intermittent “flashing” artifacts in fUS images that may bias the ΔCBV estimates. We implemented temporal filtering and intensity thresholding to reduce these artifacts, but future studies may be needed to investigate and find solutions to systematically remove these flashing artifacts. Second, ULM can be influenced by variabilities in microbubble injection and physiological conditions of the animal. In this study we used fixed total number microbubble counts to retrospectively control these variations but a future study may be needed to develop a prospective and adaptive microbubble concentration control method to acquire ULM data with consistent microbubble distributions in the blood stream. Third, although head fixation minimized large tissue displacements in our study, small, micron-level movements were still present in our dataset as manifested by the blurring artifacts in the ULM images. Lastly, although imaging in the awake state preserves native neurovascular dynamics, it unavoidably introduces behavioral variability. For instance, whisker-evoked responses are susceptible to modulation by stress, arousal, or anxiety, particularly during early habituation phases. Further refinement of the awake imaging paradigms, including freely moving approaches ^27^ or roller-supported head-fixation setups ^28^, may help reduce anxiety-related confounds while preserving physiological brain states.

In summary, in this paper we developed a multiscale and multimodal ultrasound imaging framework that integrates brain-wide fUS with microvessel-scale ULM and fULM in awake mice. Across five months of longitudinal measurements, fUS reliably mapped whisker-evoked hemodynamic activation, while ULM and fULM provided complementary microvascular structural and functional readouts. Importantly, the framework supported prolonged longitudinal imaging without evidence of chronic neuroinflammation or memory impairment, and the animals remained healthy throughout the study. Together, these results establish practical baseline variability and sensitivity benchmarks for future longitudinal studies using these ultrasound technologies and provide a broadly applicable tool for neuroimaging research in awake mice.

## 4 Methods

### Animal preparation

Only male 5xFAD-negative non-transgenic littermate mice were used in this study as healthy controls. These mice were 12 weeks old at the time of surgery. Male mice were used to reduce potential biological variability associated with sex-dependent vascular and physiological fluctuations, including those related to the estrus cycle ^17^. The 5xFAD colony was maintained by crossing B6SJL-Tg(APPSwFlLon,PSEN1M146LL286V)6799Vas/Mmjax transgenic mice with B6SJLF1/J mice. Genotyping was performed using the Transnetyx service, and mice negative for the 5xFAD transgene were used as non-transgenic littermate controls in the present study.

All procedures were approved by the Institutional Animal Care and Use Committee of the University of Illinois Urbana–Champaign (Protocol #22033). Mice were maintained under a 12-h light/dark cycle with ad libitum access to food and water. Animals were group-housed before surgery and singly housed after cranial window implantation to facilitate postoperative recovery and reduce the risk of injury.

### Pre-surgery handling

Beginning one week before cranial window surgery, mice were habituated by tunnel handling, a method shown to lower anxiety in mice ^29^. For the first 3 days, a commercially available polycarbonate transfer tube (TRANS-TUBE, 130 × 50 mm; Braintree Scientific, Inc., Braintree, MA, USA) was used. Mice were encouraged to voluntarily enter the tube from the home cage, and the tube was then gently raised while the animal stayed inside for approximately 30 s. The mouse was then returned to the cage for one minute of free movement before the handling procedure was repeated. This routine was carried out twice daily. After approximately three days, the mice became accustomed to the tunnel handling process. At this point, a 3D-printed body tube was attached to one end of the commercial tunnel, and the mice were allowed to enter the body tube voluntarily. For the subsequent five days, the body tube replaced the commercial tunnel for handling, ensuring that the mice were comfortable and familiar with the procedure.

### Cranial window surgery

Cranial window implantation was performed as previously described 29^14,19^.To reduce perioperative brain swelling, mice received dexamethasone (0.5 mg/kg, intraperitoneal) before surgery. Anesthesia was induced with 3% isoflurane and maintained at 1.0%–1.5% throughout the procedure. After fixation in a stereotaxic frame, a midline scalp incision was made and the temporalis muscle was retracted from the skull. A 5 mm (AP) × 8 mm (ML) cranial window was opened with a dental drill under continuous sterile saline irrigation, and the skull flap was removed while preserving the dura. A polymethylpentene (PMP) sheet (Goodfellow, ME311051) was placed over the window and secured with tissue adhesive and dental cement. A headpost was attached to the remaining skull using the same materials (Super-Bond C&B, Sun Medical Co., Ltd.). The window was then sealed with biocompatible silicone rubber (Body Double-Fast Set, Smooth-On). After the silicone cured, anesthesia was discontinued and the animal was allowed to recover.

### Post-surgical care and head-fix habituation

Postoperative analgesia was provided by subcutaneous carprofen (5–10 mg/kg) immediately after surgery. Mice were placed in individual cages post-surgery and monitored every 15 minutes until regaining sternal recumbency. If signs of discomfort were observed, additional doses of Carprofen (5-10 mg/kg) were administered subcutaneously every 24 hours, as needed, for up to five days. Animals were monitored daily for two weeks to ensure proper healing. Following recovery, mice underwent daily head-fixation habituation, beginning with voluntary entry into a 3D-printed body tube. Head-fixation duration was progressively increased from 10 min on day 1 to 1 h by day ^14^. Sessions were immediately stopped if animals exhibited signs of distress, including excessive struggling or vocalization.

### Experimental procedure of imaging sessions

On each imaging day, mice were guided into the body tube, which was fixed to the imaging table, and head fixation was achieved by securing the headpost to the tube. The protective silicone layer was removed from the cranial window using forceps, after which ultrasound coupling gel was applied and the transducer was positioned to identify the imaging plane. During the first session, the target coronal plane was defined at bregma −1.5 mm using anatomical landmarks. Once identified, the relative position of the probe and headpost was recorded to ensure consistent repositioning in subsequent sessions. B-mode and power Doppler images were also saved to assist relocation of the same imaging plane over time.

Imaging was performed using a Vantage 256 ultrasound system (Verasonics Inc., Kirkland, WA) equipped with an L22--14vX linear-array transducer (Verasonics Inc.). The probe was mounted in a custom 3D-printed holder attached to a motorized translation stage (VT-80, Physik Instrumente, Auburn, MA), enabling precise adjustment in the elevational direction to target the desired coronal plane. For each mouse, imaging was performed in a fixed sequence of task-evoked fUS, ULM, and task-evoked fULM.

#### Task-evoked fUS

To minimize motion artifacts, mice were head-fixed and acclimated for at least 5 minutes before data collection. Imaging was performed in an awake state under low-light conditions to reduce external stimuli that might interfere with spontaneous neural activity. Whisker stimulation was delivered using a custom-built device, an Arduino-controlled wooden dowel oscillated at a rate of 3 Hz within a fixed angular range, to deflect the mouse’s whiskers. The wooden dowel was positioned 5-7 mm from the targeted mystacial (whisker) pad on the face, oriented perpendicular to the whiskers to produce deflection without skin contact. Every time the stimulation data acquisition started, the Arduino received an output trigger from the Verasonics system, and then initiated a period of whisker stimulation by driving the wooden dowel. Each trial lasted a total of 360 seconds, including 30 seconds of baseline recording, 5 on-off blocks (30 seconds of stimulation, 30 seconds of rest in each block), and 30 seconds of post-stimulation rest. Ultrasound was transmitted at a center frequency of 15.625 MHz with a 5-angle compounding sequence (−2°, −1°, 0°, 1°, and 2°) that results in a post-compounding frame rate of 500 Hz. Each acquisition used a Doppler ensemble size of 250 frames, which were beamformed in real-time using the Verasonics reconstruction program. The final fUS imaging frame rate was established at 1Hz for functional imaging.

#### ULM

Prior to tail vein cannulation, mice were briefly anesthetized with isoflurane (3% induction, 1% maintenance) to minimize pain and stress. A 29-gauge catheter was inserted into the tail vein of each mouse and anesthesia was subsequently discontinued. The animals were allowed to recover and regain full movement of their limbs and tail. A programmable infusion pump (NE-300, New Era Pump Systems Inc., Farmingdale, NY) was used to administer fresh, activated Lumason^®^ (Bracco Diagnostics Inc., Monroe Township, NJ, USA) at a rate of 30 μl/min through the tail vein catheter. MB solutions were mixed every 5 minutes using a magnetic stirrer to maintain a constant MB concentration during the experiment. The same imaging sequence for fUS described above was used for ULM and fULM, except that fUS data were acquired at a transmit voltage of 40 V, whereas ULM and fULM data were acquired at 6 V. Instead of real-time beamforming, for ULM, we elected to continuously acquire 800 imaging frames per second (in the format of raw, unbeamformed RF channel data) to facilitate robust ULM reconstruction. Offline beamforming was subsequently performed using the Verasonics reconstruction program after completion of the experiment. Task-evoked fULM

The procedure involved administering fresh, activated Lumason^®^ through the tail vein catheter at a rate of 60 μl/min. The whisker stimulation paradigm was identical to that used in the fUS experiments (5 on-off blocks, 360s in total). The data collection procedure followed the ULM acquisition, which continuously saved 800 frames of RF data per second (1000 Hz PRF) for subsequent offline beamforming.

### Processing of images and image analysis

#### 1. fUS

Raw ultrasound data were processed using a spatiotemporal clutter filter based on singular value decomposition (SVD) to suppress tissue motion and background noise ^30^. The first 50 singular values, primarily representing tissue-related components, were discarded. The remaining signals were averaged over time to generate power Doppler images that served as CBV reference maps. For each session, we then derived two functional maps from the CBV time series: a *z*-score activation map, which captures the statistical significance of stimulus evoked responses, and a ΔCBV map, which quantifies the response magnitude. The *z*-score was calculated according to Fisher’s transform with the following formula: 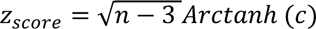, where n denotes the total number of time samples in the session (e.g., n = 360 in this case), and *c* represents the Pearson’s correlation coefficient between the time series of each pixel (360 time points) and the stimulation pattern ^19^. A Bonferroni-corrected significance threshold of *p* < 0.05 was applied to identify activated regions ^31^. Finally, the *z*-score was overlaid on grayscale power Doppler images for visualization. ΔCBV was computed as a percent change relative a block-specific pre-stimulus baseline. Specifically, within each stimulus block, all samples from the 15-s pre-stimulus window, the subsequent 30-s stimulation, and the 15-s post-stimulation rest were normalized to the mean CBV measured during the 15 s immediately preceding that block’s stimulus onset. We refer to the concatenated 15-s pre-stimulus, 30-s stimulation, and 15-s post-stimulation rest within each block as a “stimulation cycle.” A ΔCBV map was then generated by averaging the ΔCBV values across the 30-s stimulation period within each cycle. For visualization of the ΔCBV maps, a correlation-based mask was applied to suppress low-signal regions. Specifically, only pixels with a correlation value greater than 0.3 were retained, and ΔCBV was displayed within this masked region. To enable anatomical quantification, each imaging was manually aligned to the corresponding anatomical plane of the Allen Mouse Brain Atlas based on visible structural landmarks, and regions of interest (ROIs) were delineated according to atlas-defined boundaries. The mean ΔCBV within ROI was then computed for subsequent statistical analysis.

### 2. ULM and fULM

SVD filtering was used to extract MB signals from the tissue background in each IQ dataset. An adaptive SVD threshold was chosen to better accommodate variable tissue motion in the experimental data ^32,33^. A 2D Gaussian function was fitted to the axial and lateral dimensions of manually selected MBs to represent the system’s Point Spread Function (PSF). After applying SVD filtering, MB separation ^34^ was performed on the IQ data, followed by 2D spline interpolation ^35^ to an isotropic resolution of 4.928 µm in both axial and lateral dimensions. The interpolated dataset underwent normalized 2D cross-correlation with the empirical PSF to identify candidate MB locations. Pixels with low cross-correlation coefficients were excluded ^34^, and regional maxima in the resulting cross-correlation map were used to identify the centroid locations of individual MBs ^36^. Tracking and trajectory estimation of MB centroids were carried out using the uTrack algorithm ^37^, with a minimum trajectory length of 10 frames (10 ms) applied for the super-resolution reconstruction. To compensate for inter-session spatial offsets during longitudinal accumulation, a post hoc rigid in-plane motion correction based on normalized cross-correlation was applied to ULM maps. Translational offsets in the axial and lateral directions were estimated from the microbubble density maps and used to realign all ULM images by linear interpolation before accumulation; acquisitions with estimated displacements greater than 100 pixels in either the axial or lateral direction were excluded. The accumulated data from each acquisition session were used to generate the final ULM vascular reconstruction of the cerebral vasculature. ROIs were delineated according to atlas-defined anatomical boundaries. Brain vascularity within each ROI was calculated by binarizing the super-resolution vessel maps and determining the percentage of area containing MB-localized pixels. Blood vessel velocity was determined for every MB track directly from the frame-to-frame displacement of detected MB centroids.

#### Stimulus-evoked fULM

For fULM, the pre-processed IQ datasets underwent the same SVD-based clutter filtering, MB localization, and tracking procedures as described for ULM. In addition to vascular reconstruction, the temporal dynamics of MB counts were extracted at each pixel to capture hemodynamic responses. Specifically, the number of localized MBs was accumulated in one-second frame over the duration of each trial, resulting in a MB count time series. For each pixel, the MB count trace was correlated with the stimulation pattern using Pearson’s correlation analysis, yielding a correlation coefficient map. Finally, the functional correlation maps were registered to the atlas for quantitative analysis within specific brain regions.

### Histological analysis

Brain tissue was harvested and fixed in 4% paraformaldehyde in PBS at 4°C for 24-48 hours, then cryoprotected in 10%, 20%, and 30% sucrose solutions. Fixed brains were embedded and sectioned at 20 μm thickness using a cryostat (CM1850, Leica, NJ, USA). Sections were washed in TBS and permeabilized with 0.3% Triton X-100 (Sigma-Aldrich, St. Louis, MO, USA), then blocked with 0.3% Triton X-100, 5% normal goat serum (Sigma-Aldrich, St. Louis, MO, USA), and 3% BSA (Sigma-Aldrich, St. Louis, MO, USA). Primary antibodies-Rabbit Anti-GFAP (Abcam, ab7260; 1:1000) and Goat Anti-Iba1 (Novus, NB100-1028; 1:1000)-were applied overnight at 4°C, followed by Alexa Fluor 488-conjugated anti-rabbit IgG (Invitrogen, A-11008; 1:1000) and Alexa Fluor 594-conjugated anti-goat IgG (Invitrogen, A-11058; 1:1000) for 2 hours at room temperature. After final washes, sections were mounted with VECTASHIELD Antifade Mounting Medium with DAPI (Vector Laboratories, Burlingame, CA, USA) and imaged using a Zeiss LSM 710 confocal microscope. GFAP, Iba1, and DAPI signals were acquired using 488 nm, 561 nm, and 358 nm excitation lasers, respectively, under identical acquisition settings. Maximum intensity projections were generated using Zen Blue. ROIs were manually defined based on DAPI morphology, and fluorescence intensities for GFAP and Iba1 were quantified in ImageJ with background subtraction.

### Passive Avoidance Test

The passive avoidance test was conducted using a two-compartment apparatus consisting of an illuminated chamber and a dark chamber equipped with a grid floor for foot shock delivery ^38^. Mice were trained once daily from D1 to D7. During each training session, the mouse was placed in the illuminated chamber, and the step-through latency to enter the dark chamber was recorded. Upon entry, a brief foot shock was delivered. Training latency was capped at 120 s. On D8, mice underwent the retention test, during which step-through latency was measured again as an index of memory retention, with a maximum cutoff time of 300 s.

### Statistical analysis

Statistical analyses were performed using GraphPad Prism (version 9.1.2, GraphPad Software, CA, USA) and Python (version 3.9, Python Software Foundation, USA). For stimulus-evoked hemodynamic responses measured by fUS and fULM, ΔCBV and ΔMB flow (%) time courses were averaged across five repeated stimulation trials within each imaging session to generate one session-level mean value per animal. These session-level summary values were used for all subsequent analyses. For longitudinal imaging analyses, n = 7 biologically independent mice were included, each measured repeatedly across five monthly sessions. Individual animals were treated as biological replicates, whereas the five repeated stimulation trials within each imaging session were treated as repeated measurements rather than independent replicates. To assess month-to-month differences in the figure panels, session-level values for fUS ΔCBV, vascularity, mean flow velocity, fULM Δ CBV, and fULM flow-stimulus correlation were compared across the five monthly sessions using one-way repeated-measures analysis of variance (ANOVA). All tests were two-sided, and a p value < 0.05 was considered statistically significant. To evaluate long-term stability, paired two one-sided tests (TOST) were performed to compare Month 3 and Month 7 measurements. Equivalence bounds were defined for each metric using the MDC95. Statistical equivalence was concluded when the 90% confidence interval of the paired difference was fully contained within the corresponding equivalence margin, with both one-sided tests significant at α = 0.05.

To further assess longitudinal stability and test for systematic time-dependent bias while accounting for repeated measurements within subjects, the same session-level values were additionally analyzed using linear mixed-effects models, with session included as a fixed effect and subject as a random effect. The p values derived from these mixed-effects models are reported in Table 1 as trend p values. To quantify longitudinal baseline variability and reproducibility, within-subject CV, ICC, RC, reference intervals, and MDC95 were calculated from repeated measurements across sessions. Within-subject CV was calculated separately for each mouse as the standard deviation across longitudinal sessions divided by the corresponding mean, multiplied by 100%. The value reported in Table 1 represents the mean of these mouse-specific CV values across animals. The RC was calculated as 1.96 × √2 × 𝑆_w_, where 𝑆_𝑤 is the within-subject standard deviation. The minimal detectable change at the 95% confidence level (MDC95) was calculated as 𝑀𝐷𝐶_95_ = 1.96 × √2 × 𝑆𝐸𝑀. Under the definitions used here, RC and MDC95 were numerically identical because they were derived from the same within-subject error term.

## Author contributions

Conceptualization and Funding Acquisition: P. Song, D. A. Llano; Methodology: P. Song, Z. Huang, Y. Wang, M. R. Lowerison, Y. Xu, Y. Shin, and B.-Z. Lin; Software: P. Song, Y. Wang and M. R. Lowerison; Validation: Z. Huang and Y. Wang; Formal Analysis: Z. Huang; Investigation: Z. Huang and Y. Wang; Resources: B.-Z. Lin, N. V. C. Sekaran, D. A. Llano, and P. Song; Data Curation: Z. Huang; Writing - Original Draft: Z. Huang, Y. Wang, P. Song; Writing - Review & Editing: All authors; Visualization: Z. Huang, Y. Wang, Y. Xu, and Y. Shin; Project Administration: P. Song. All authors have read and approved the final version of the manuscript.

## Materials & Correspondence

Pengfei Song

## Acknowledgements

This study was partially supported by the National Institute of Biomedical Imaging and Bioengineering and the National Institute of Neurological Disorders and Stroke of the National Institutes of Health under grant numbers R21EB030072 and R56NS131516, by the National Science Foundation CAREER Award 2237166, and by the Chan Zuckerberg Initiative Ben Barres Early Career Award. The content is solely the responsibility of the authors and does not necessarily represent the official views of the NIH and NSF.

## Abbreviations

CBF: cerebral blood flow
fUS: functional ultrasound
ULM: ultrasound localization microscopy
fULM: functional ultrasound localization microscopy
CV: coefficient of variation
ICC: intraclass correlation coefficient
RC: repeatability coefficient
MDC95: minimal detectable change at the 95% confidence level
SVD: singular value decomposition
GLM: general linear model
ROI: region of interest
PSF: point spread function
IQ: in-phase quadrature.

---

**Supplementary Fig. 1.**
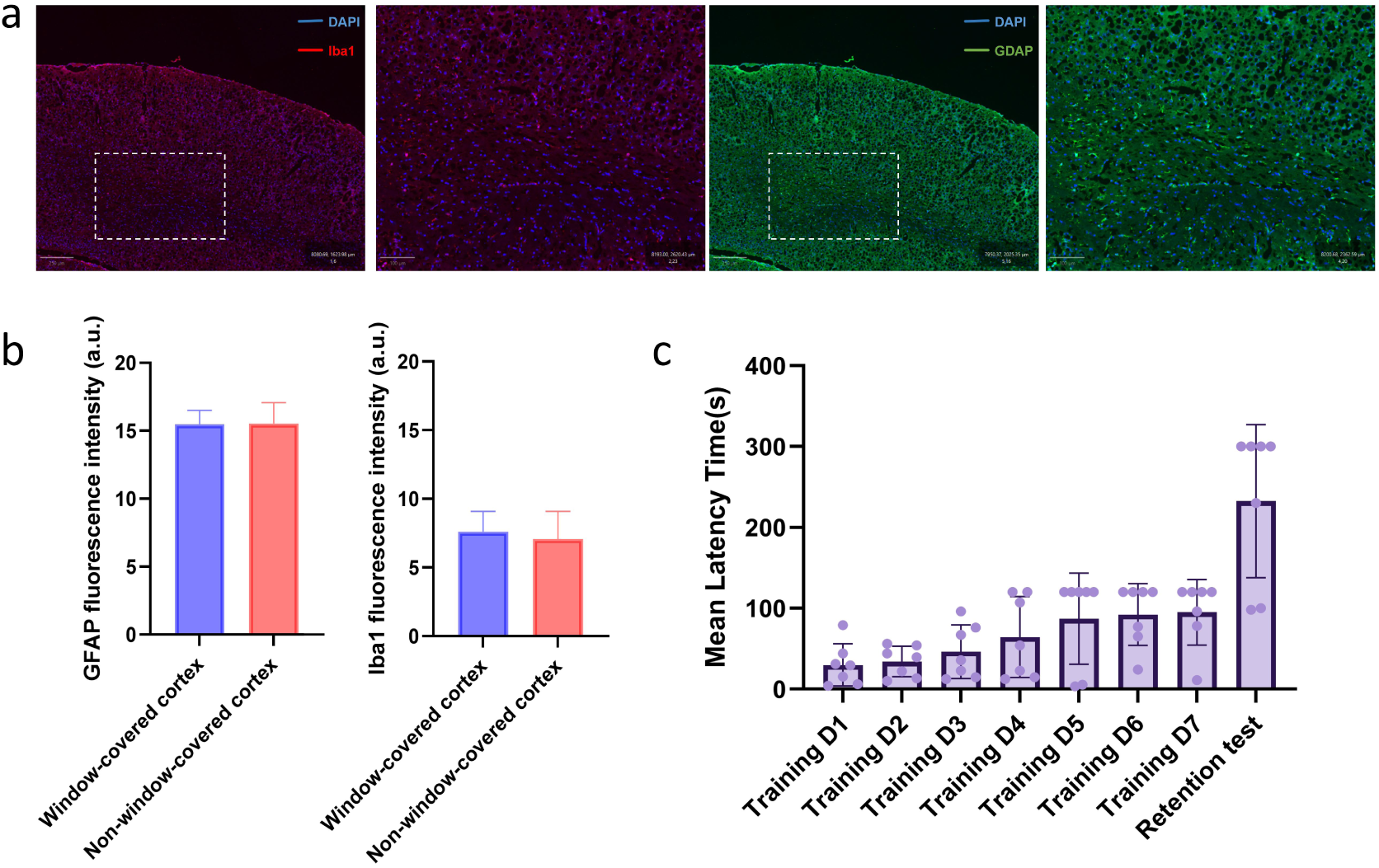
Histological and behavioral assessment of long-term cortical biocompatibility after chronic PMP cranial window implantation. (a) Representative confocal images of cortical sections collected after five months of repeated awake imaging. Iba1-positive microglia are shown in red, GFAP-positive astrocytes in green, and nuclei (DAPI) in blue. No obvious reactive gliosis or focal inflammatory cell accumulation was observed beneath the chronic PMP window. Scale bars, 200 μm. (b) Background-subtracted fluorescence intensities of GFAP and Iba1 in the window-covered cortex and non-window-covered cortex. (c) Passive avoidance testing showed prolonged step-through latency during retention testing. Data are presented as mean ± SD.

